# Utilisation of semiconductor sequencing for the detection of predictive biomarkers in glioblastoma

**DOI:** 10.1101/2021.01.11.426191

**Authors:** Gareth H Williams, Robert Thatcher, Keeda-Marie Hardisty, Marco Loddo

## Abstract

The standard treatment for glioblastoma involves a combination of surgery, radiation and chemotherapy but have limited impact on survival. The exponential increase in targeted agents directed at pivotal oncogenic pathways now provide new therapeutic opportunities for this tumour type. However, lack of comprehensive precision oncology testing at diagnosis means such therapeutic opportunities are potentially overlooked.

To investigate the role of semiconductor sequencing for detection of predictive biomarkers in routine glioblastoma samples we have undertaken analysis of test trending data generated by a clinically validated next generation sequencing platform designed to capture 764 of the leading anti-cancer targeted agents/combinations and immunotherapies via analysis of actionable genomic variants distributed across 505 genes. Analysis was performed across a cohort of 55 glioblastoma patients.

Analysis of trending data has revealed a complex and rich actionable mutational landscape in which 166 actionable mutations were detected across 36 genes linked to 17 off label targeted therapy protocols and 111 clinical trials. The majority of patients harboured three or more actionable mutations affecting key cancer related regulatory networks including the PI3K/AKT/MTOR and RAS/RAF/MEK/MAPK signalling pathways, DNA-damage repair pathways and cell cycle checkpoints. Linkage with immunotherapy and PARP inhibitors was identified in 44% of glioblastoma patients as a consequence of alterations in DNA-damage repair genes.

Taken together our data indicates that precision oncology testing utilising semiconductor sequencing can be used to identify a broad therapeutic armamentarium of targeted therapies and immunotherapies that can be potentially employed for the improved clinical management of glioblastoma patients.

## Introduction

Gliomas are the most common tumors of the primary central nervous system, and of these, over a half of cases represent glioblastoma (GBM) at diagnosis. In the US approximately 70,000 primary CNS tumors are diagnosed, GBM being the most frequent high-grade glioma, with an incidence of 3–4/100,000 (1). This has the most malignant phenotype of primary brain cancers and is associated with a poor prognosis with a median survival of around 14-18 months (2). Its inherently disabling effects on patients, often to prevent independent function, result in a significant burden on health-care systems which is out of proportion with its incidence. The present conventional treatment protocols (surgery, radiation and the alkylating chemotherapy agent temozolomide) have limited impact in improving patient survival (3–5). A range of new approaches have been developed to broaden the potential therapies available to glioblastoma patients, which include the use of Gamma Knife radiosurgery, hyperthermia and the development of new pharmacological tools such as targeted therapies and immunotherapies (6).

The last decade has seen major advances in dissecting the aberrant molecular pathways that contribute to glioblastoma development. Comprehensive genetic screens have shown that genetic alterations in glioblastoma are distributed across the entire genome, resulting in the dysregulation of many critical signaling pathways such as the RB and p53 pathways and the receptor tyrosine kinase/Ras/phosphoinositide 3-kinase signaling pathways (7, 8). There are also errors in DNA replication, DNA repair, chromosomal segregation and disruption of cell cycle checkpoints which all contribute to uncontrolled cell proliferation (9–11). The wide range of molecular events in this tumor type therefore provides new potential therapeutic opportunities using the new generation of targeted anti-cancer agents including small molecule inhibitors and humanized monoclonal antibodies which have been specifically developed against these molecular targets.

The increasing use of targeted agents offers the great advantage of increased specificity and reduced toxicity when compared with conventional chemotherapy (12). Meta-analysis in diverse tumor types has shown that a personalized strategy of treatment is an independent predictor of better outcomes and fewer toxic deaths when compared with chemotherapy (13). However, it is critical that targeted agents are precisely matched to the correct patients based on their tumor mutation profile. When prescribed in the absence of tumor molecular profiling, targeted agents actually have poorer outcomes when compared with cytotoxics (13). Identification of the most appropriate targeted therapies for cancer patients is currently a challenge because molecular profiling tests used in routine clinical practice are limited in their coverage and do not provide direct evidence based-linkage with therapy. Moreover, many sequencing platforms are unable to operate efficiently when applied to small clinical biopsy samples which have undergone formalin fixation resulting in low DNA/RNA yields with low integrity and quality (14). At the present time routine molecular testing with regard to glioblastoma is restricted to a limited number of biomarkers such as IDH mutation status, MGMT promoter methylation and 1p19q co-deletion FISH analysis (15). These biomarkers provide prognostic information and weak prediction of chemotherapy response. However, the new era of targeted next-generation sequencing (NGS) technologies, optimized for nucleic acid templates extracted from routine formalin fixed, paraffin embedded tumor samples (PWET), now provides the opportunity for comprehensive predictive testing in solid tumors.

To investigate the potential role of clinically directed semiconductor targeted sequencing in solid tumours we have established a clinically validated next generation sequencing (NGS) platform optimized for the analysis of PWET clinical biopsy samples. This platform enables capture of 764 lead anti-cancer targeted agents/combinations and immunotherapy opportunities via analysis of actionable variants distributed across 505 genes. All actionable variants analysed are linked to targeted therapies either on-market FDA and EMA approved, carrying ESMO and NCCN guideline references or currently in clinical trials, phases I-IV, worldwide

Here we have conducted a retrospective analysis of precision oncology profiling test trending data relating to a cohort of 55 glioblastoma patients who underwent testing following failure of first line treatment protocols. This analysis has shown that semiconductor sequencing can be applied robustly to routine processed glioblastoma samples enabling detection of a broad range of actionable mutations and their cognate therapies. The high frequency of actionable mutations detected in this tumour type demonstrates the broad range of potential therapeutic opportunities that are now available to glioblastoma patients following clinically directed precision oncology profiling.

## Materials and Methods

### Patient demographics

A retrospective analysis was performed on the trending data generated as part of routine comprehensive precision oncology NGS testing for solid tumours and collected in compliance with ISO15189:2012 requirements for monitoring of quality indicators. Anonymized and coded patient data was collected between January 2018 and July 2019. A database search identified 55 cases of glioblastoma which were included in the analysis. The patient demographics including prognostic biomarker status for this cohort are shown in S1 Table. The research conducted in this study was limited to secondary use of information previously collected in the course of normal care (without an intention to use it for research at the time of collection) and therefore does not require REC review. The patients and service users were not identifiable to the research team carrying out trend data analysis. This is also in accordance with guidance issued by the National Research Ethics Service, Health Research Authority, NHS and follows the tenants of the Declaration of Helsinki.

### Comprehensive NGS genomic profiling

The NGS platform utilized for clinical testing is validated for clinical use and accredited by CLIA (ID 99D2170813) and by UKAS (9376) in compliance with ISO15189:2012 and following the guidelines published by the Association for Molecular Pathology and College of American Pathologists and IQN-Path ASBL (16, 17). The performance characteristics are shown in S2 Table. The NGS platform targets 505 genes and detects actionable genetic variants linked to 764 anti-cancer targeted therapies/therapy combinations. This includes analysis of 51 driver and 349 partner genes for detection of 867 actionable fusion genes. Genomic regions selected for analysis of clinically relevant actionable variants are shown in S3 Table.

### DNA and RNA extraction, library preparation and sequencing

DNA and RNA was extracted from PWET curls cut at 10μm or from 5μm sections mounted onto unstained glass slides using the RecoverAllTM extraction kit (Ambion, Cat no.A26069). RNA samples were diluted to 5ng/μl and reverse transcribed to cDNA in a 96 well plate using the Superscript Vilo cDNA synthesis kit (CAT 11754250). Library construction, template preparation, template enrichment and sequencing were performed using Ion AmpliseqTM library 2.0 (Cat:4480441) and the Ion 540TM OT2 kit (Cat: A27753) according to the manufacturer’s instructions. Sequencing was performed using the Ion S5 system 20 (Cat: A27212) utilising Ion 540TM chips (Cat:30 A27766).

### Quality control metrics

Sequencing runs were quality controlled using the following parameters according to the manufacturer’s instructions (Ion Ampliseq library 2.0): chip loading >60% with >45 million reads observed, enrichment 98-100%, polyclonal percentage <55%, low quality <26%, usable reads > 30% and aligned bases were ≥80%, unaligned bases were <20%, mean raw accuracy was >99% and overall read length between 100-115bp for DNA and RNA. Individual DNA sample metrics were evaluated using the following parameters: number of mapped reads >4.5 million, percent reads on target >90%, average base coverage depth >1200, uniformity of amplicon (base) coverage >90%, amplicons were required to have less than 90% strand bias with >80% of amplicons reading end to end, on-target reads >85% and target base coverage at 1x, 20x, 100x and 500x >90% (S4 Table).

### Data analysis

Sequence alignment and variant calling was performed on The Torrent Suite™ Software (5.8.0). Alignment in Torrent Suite™ Software was performed using TMAP. The output BAM file was uploaded via the Ion Reporter Uploader plugin (5.8.32-1) to The Ion Reporter™ Software (5.10.1.0). Gene fusions were reported when occurring in >40 counts and meeting the threshold of assay specific internal RNA quality control with a sensitivity of 99% and PPV of 99%. Six internal expression quality controls were spiked into each sample to monitor assay performance with an acceptance cut-off of>15 reads in 5 out 6 controls [Ion Reporter™ 5.10.1.0; default fusion view (5.10). Hotspot variants with >10% alternate allele reads were classified as ‘detected’ with an assay sensitivity and positive predictive value (PPV) of 99% (S2 Table). For copy number variants (CNV), amplifications of CN> 6 with the 5% confidence value of ≥4 after normalization and deletions with 95% CI ≤1 were classified as present when the tumor content was >50% with a sensitivity of 80% and PPV of 100% (S2 Table). Variants meeting the previously defined criteria were then assessed for pathogenicity and linkage to cognate therapies using several resources including USCS, COSMIC, ClinVar databases, PhyloP, SIFT, Grantham, ExAC, DrugBank, NCI-MATCH, PFAM, DGV, MAF, GlobalData, PolyPhen in silico tools and IGV visualisation software. The results of variant annotation were organized hierarchically by gene, alteration, indication and level of evidence in relation to clinical actionability following the joint recommendation of the association of the AMP/ASCO/CAP (16) and the ESMO Scale for Clinical Actionability of Molecular Targets (18) including FDA/EMA approved therapies, guideline references by ESMO/NCCN and clinical trials (Phases I-IV) worldwide.

### Immunohistochemistry for PD-L1

An immunohistochemistry assay for the detection of PD-L1 was validated for clinical use and accredited by CLIA and by UKAS (9376) in compliance with ISO15189:2012. PD-L1 rabbit monoclonal antibody (clone E1L3N) was obtained from Cell Signalling (Cat No. 13684S). Histological sections from a representative PWET block for each case were cut at 3μm thickness and mounted on Super Frost glass slides (Leica, cat no). Section deparaffinization, antigen retrieval and immunohistochemical labelling were performed using the Bond III Autostainer and Bond Polymer Refine Detection Kit (Leica, Cat no. DS8900) according to the manufacturer’s instructions. Primary antibody was applied for 20 minutes at 1/400 dilution. Assessment of PD-L1 immunostaining was performed by a qualified histopathologist in accordance with PD-L1 clinical reporting guidelines (19).

### Statistical analysis

The 2 tailed t-test assuming equal variances was used to compare the mean number of mutations across variant type using the 95% confidence interval and assessed as statistically significant at the 5% level. All tests including means and medians were calculated using Microsoft Excel 2010.

## Results

Actionable somatic variants were identified in all glioblastoma samples tested and, more than one variant was detected in 45 out of the 55 samples (82%). Across all 55 glioblastoma samples, one hundred and sixty-six actionable mutations were detected in 36 genes (Fig 1A and S5-S7 Tables) with TP53, CDKN2A, EGFR, PTEN, CDKN2B, NF1, CDK4, IDH1, PIK3R1 and RB1 being the ten most frequently mutated (Fig 1B). The median number of actionable variants detected per sample was 3 and ranged from 1-9 (S1 Fig). Aberration of cancer genes occurs by a range of mechanisms including SNVs, gene amplification and deletion (collectively denoted CNVs), fusions and multiple nucleotide variants (MNVs) which include insertions, deletions and duplications. Their frequencies in relation to each cancer gene are shown in Figs 1B, 2 and 3. The mutational landscape is dominated by SNVs, detected in 69.1% of samples, and CNVs which occur at a similar frequency of 63.6% across all samples (S2 Fig).

**Fig 1.**
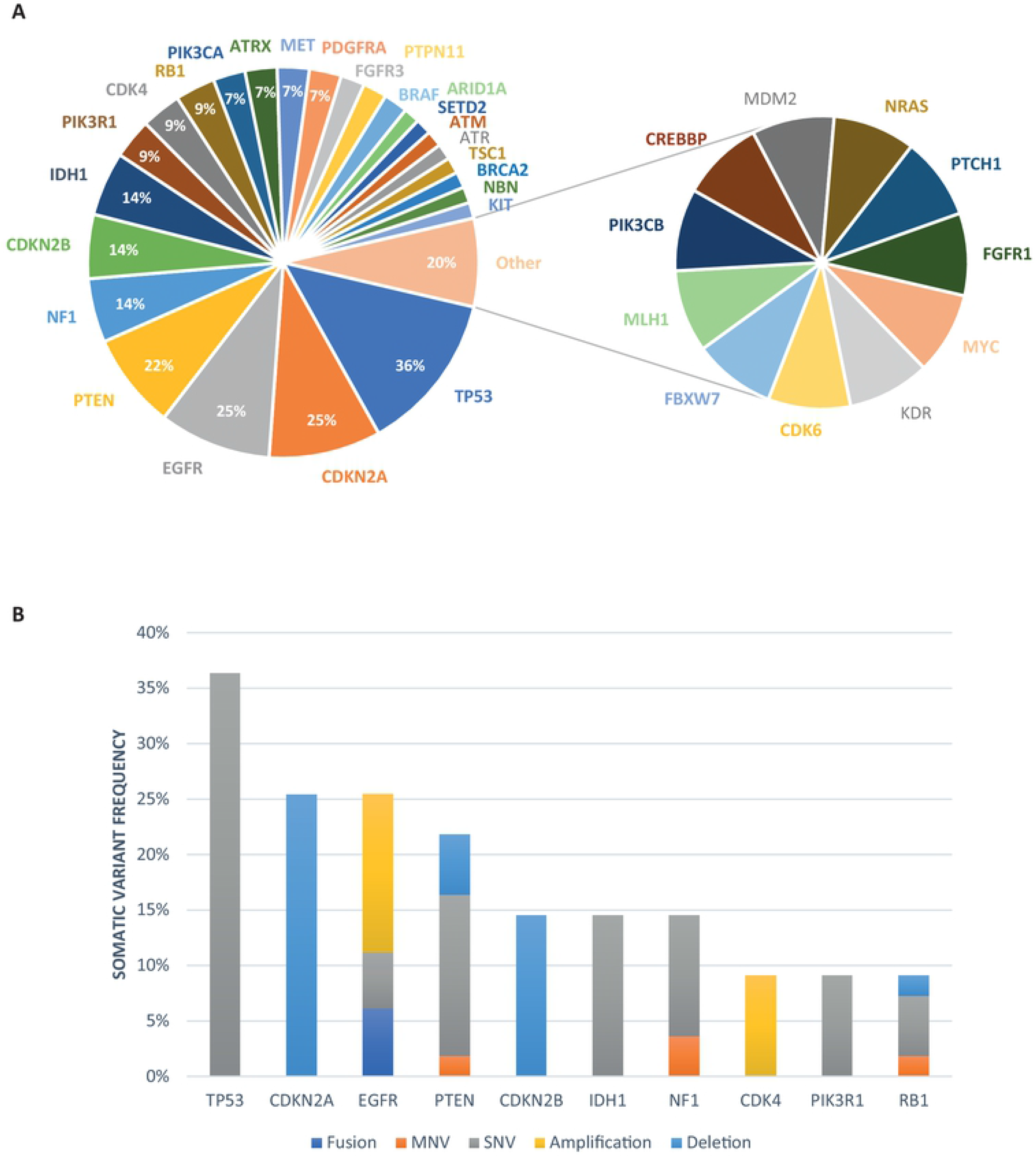
The actionable genomic landscape in glioblastoma. **A)** Pie chart showing frequency of altered genes in glioblastoma (n=55). Segments not showing percentages have a frequency of <6%; **B)** Bar chart showing then ten most frequently genetically altered genes including variant type.

**Fig 2.**
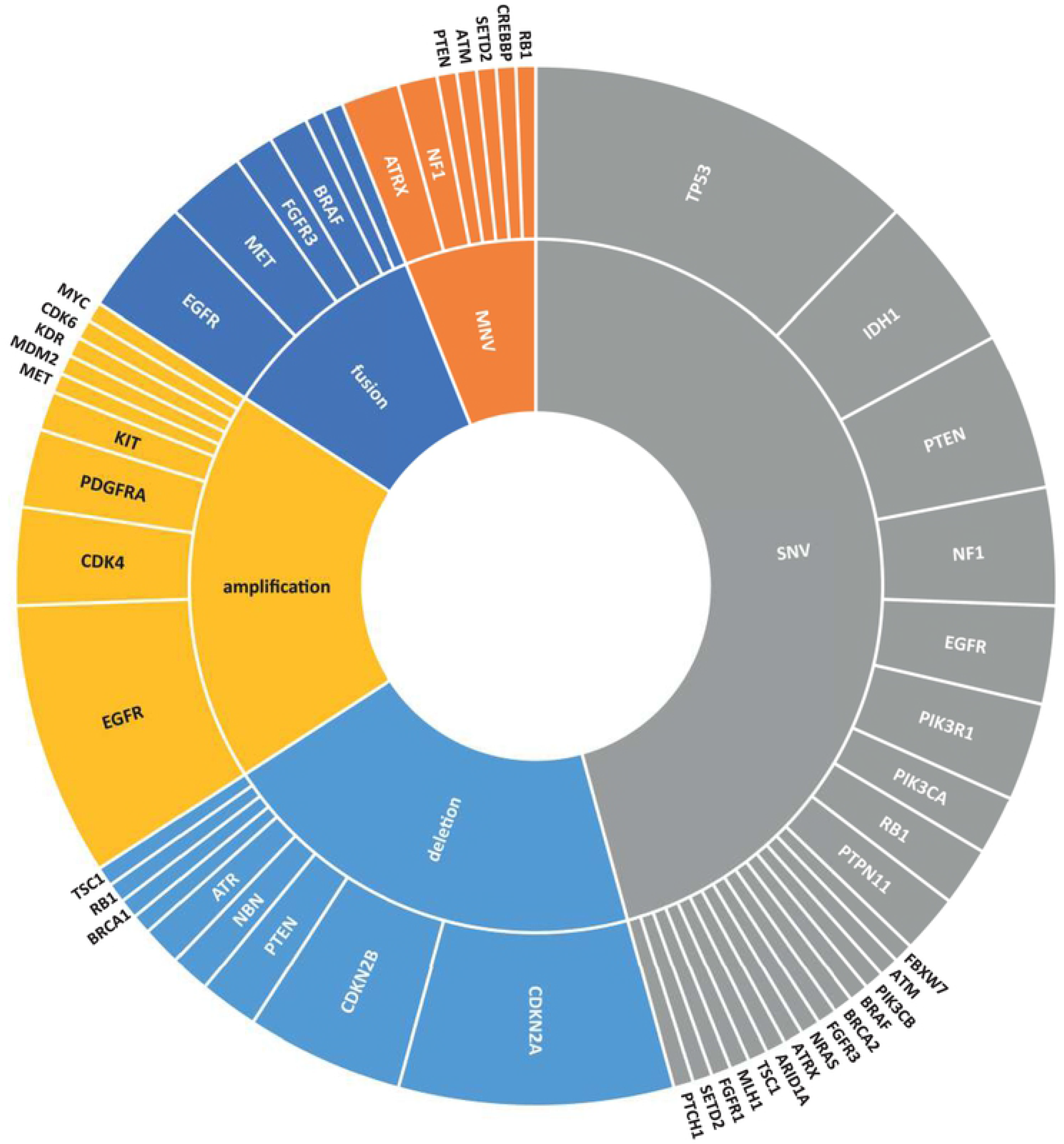
Frequency of actionable genetic alterations by variant type (inner ring) and by gene (outer ring) in all variants detected (n=164).

**Fig 3.**
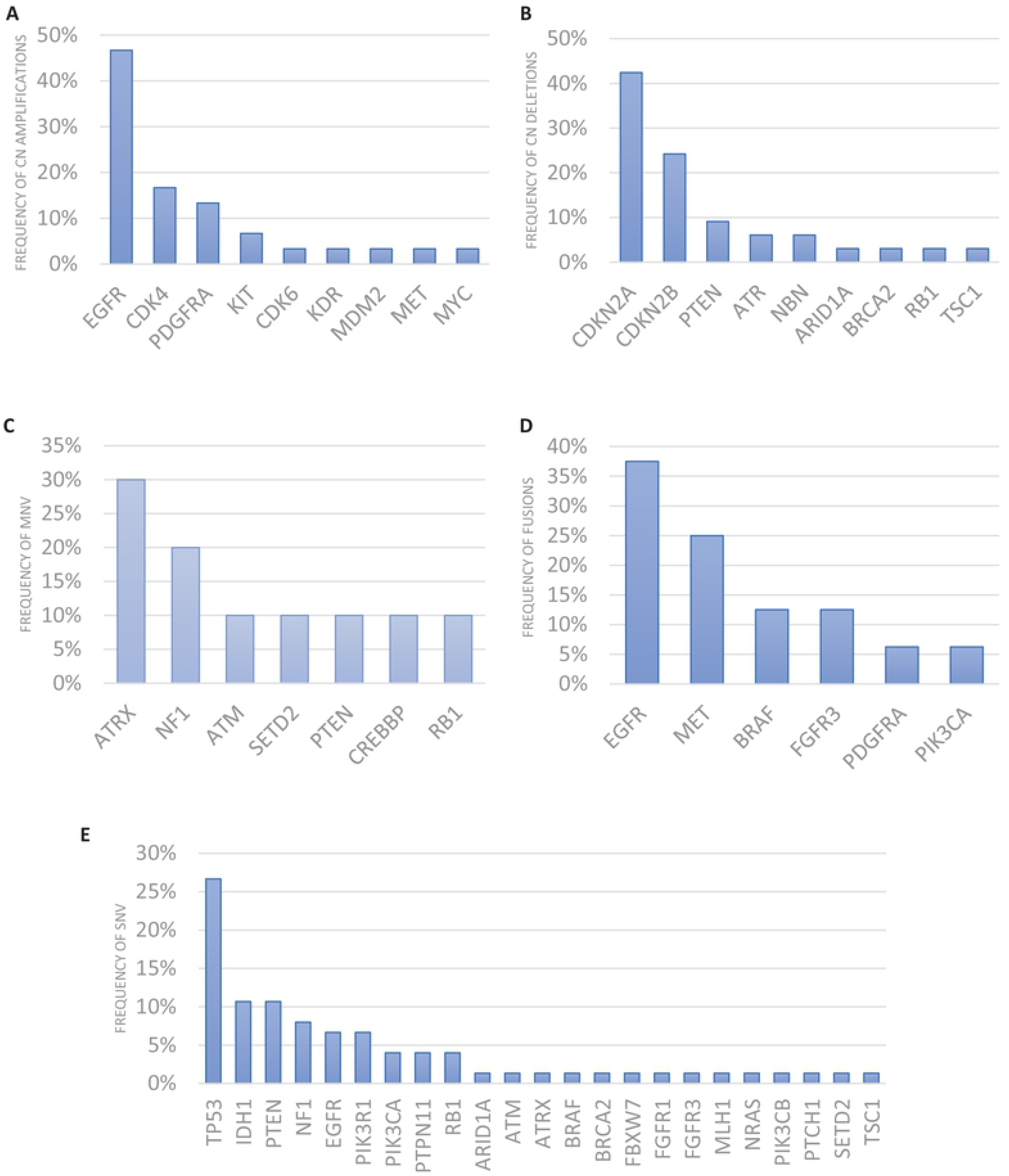
Frequency of altered genes in each variant type. **A)** Copy number amplifications; **B)** Copy number deletions; **C)** Multiple nucleotide variants (MNV); **D)** gene fusions and **E)** Single nucleotide variants (SNV).

Interestingly, a broad range of actionable fusion genes were identified in this cohort of glioblastomas including EGFR-SEPT, EFGR intragenic recombination, FGFR3-TACC3, PTPRZ1-MET, CAPZA2-MET, METex14 skipping, AGK-BRAF, EGFR Viii, TBLXR1-PIK3CA and FIP1L1-PDGFRA (Fig 3D and S7 Table). Although fusions are generally regarded as a rare event in solid tumors, with around a 3% frequency (20), here we identified druggable fusion genes in 23.6% of glioblastoma cases. Moreover, two of these fusions, TBL1XR1-PIK3CA and FIP1L1-PDGFRA, have not been previously reported in glioblastoma to our knowledge. The mean number of variants detected was significantly greater in samples containing fusions (3.92) versus absence of fusions (2.74) (p=0.025) or deletions (3.81) versus absence of deletions (2.53) (p=0.005).

All patients harbored one or more actionable variants linked bioinformatically to a wide range of targeted therapies with either FDA, EMA, NCCN or ESMO labels in relation to “cancers of other type” (off-label), or alternatively investigational targeted therapies in clinical trials (Table 1). The actionable variants in this cohort of patients were linked to 17 off-label targeted therapy protocols with FDA, EMA, NCCN or ESMO labels and meeting the tier criteria IIC level of clinical significance as defined by the Joint Consensus Recommendation of the Association for Molecular Pathology, American Society of Clinical Oncology, and College of American Pathologists (16). This cohort of patients was additionally linked to a total of 111 clinical trials (Table 1) through matching of actionable variants to their cognate targeted therapies.

**Table 1.**
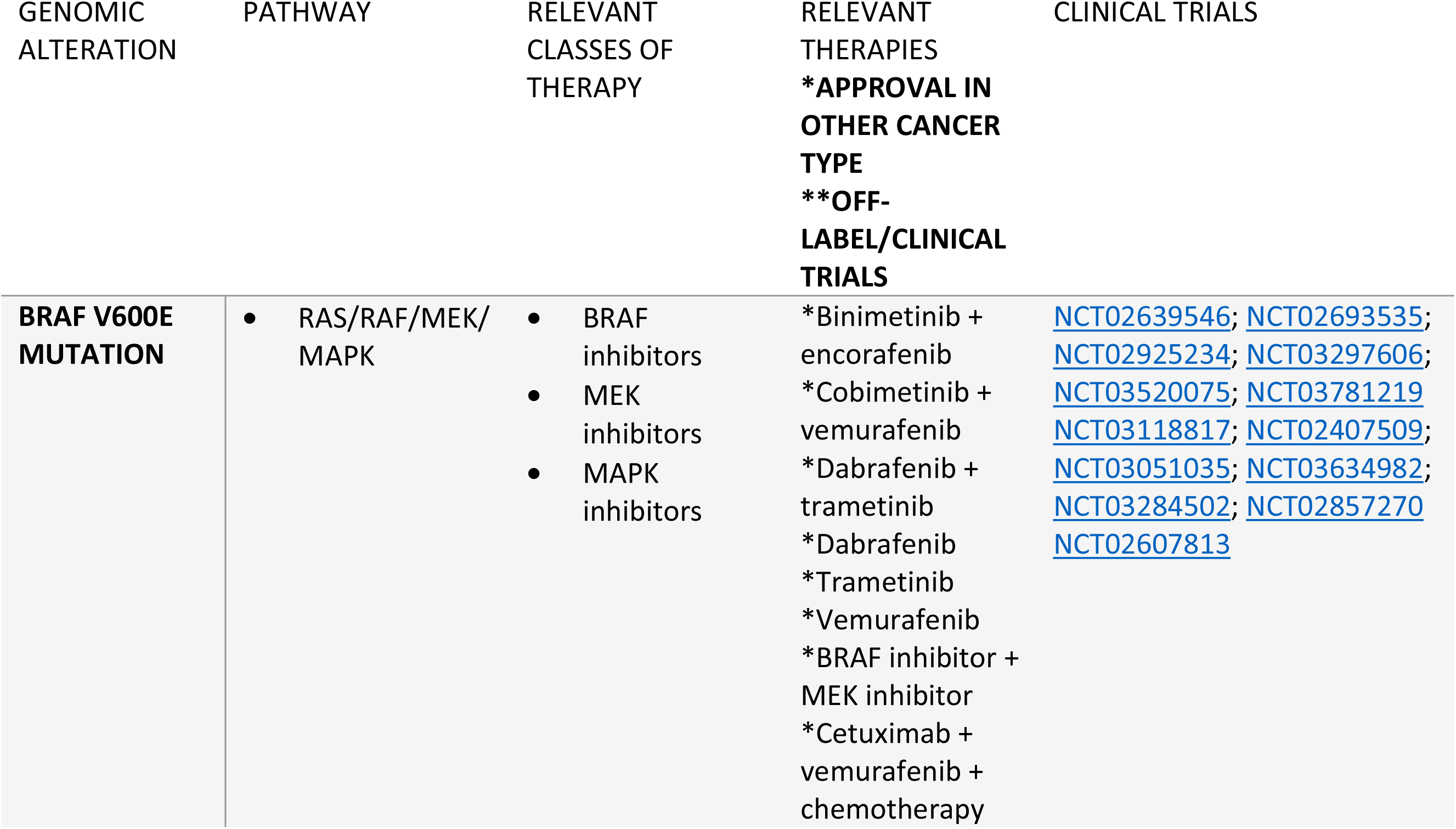
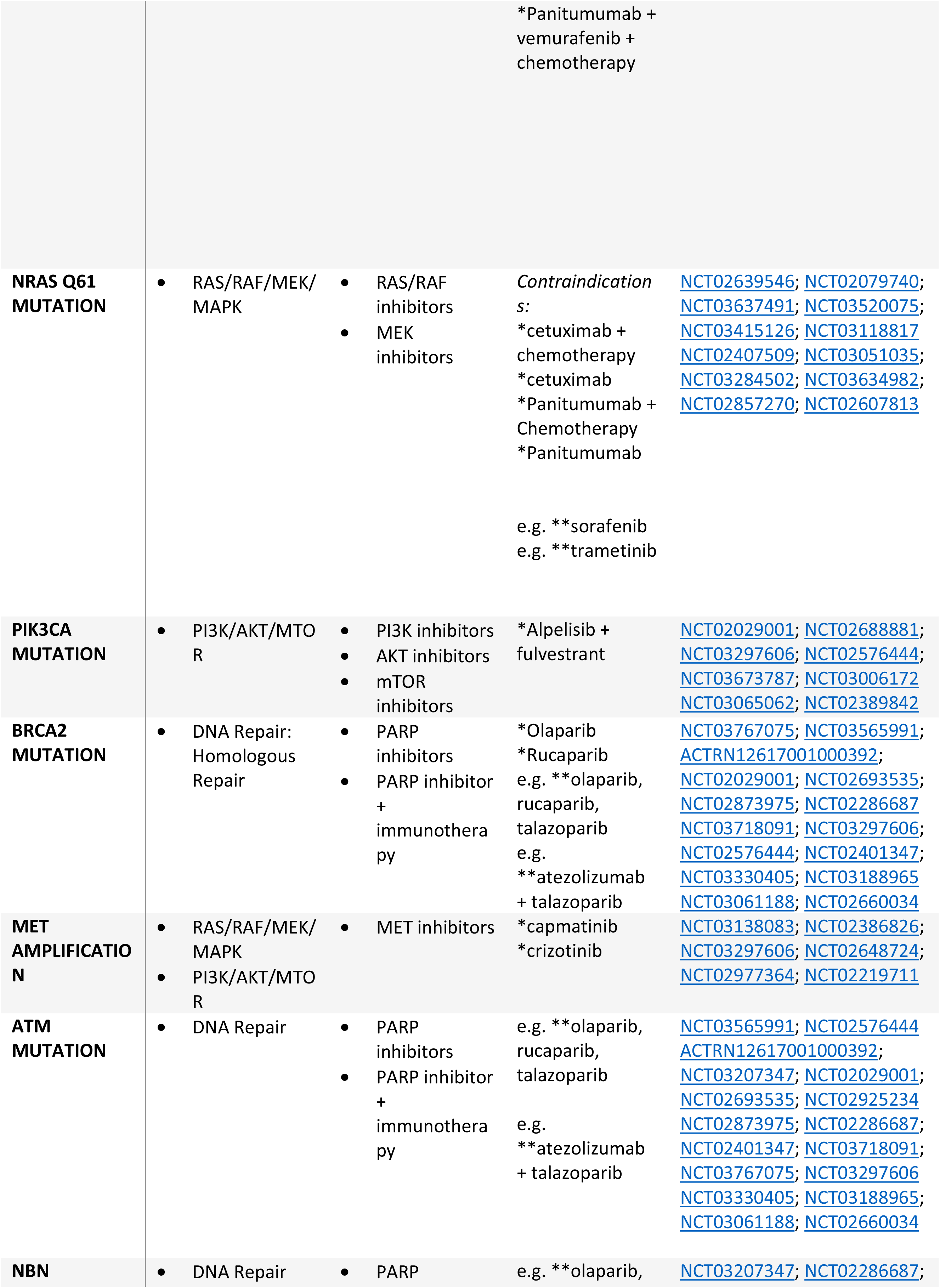
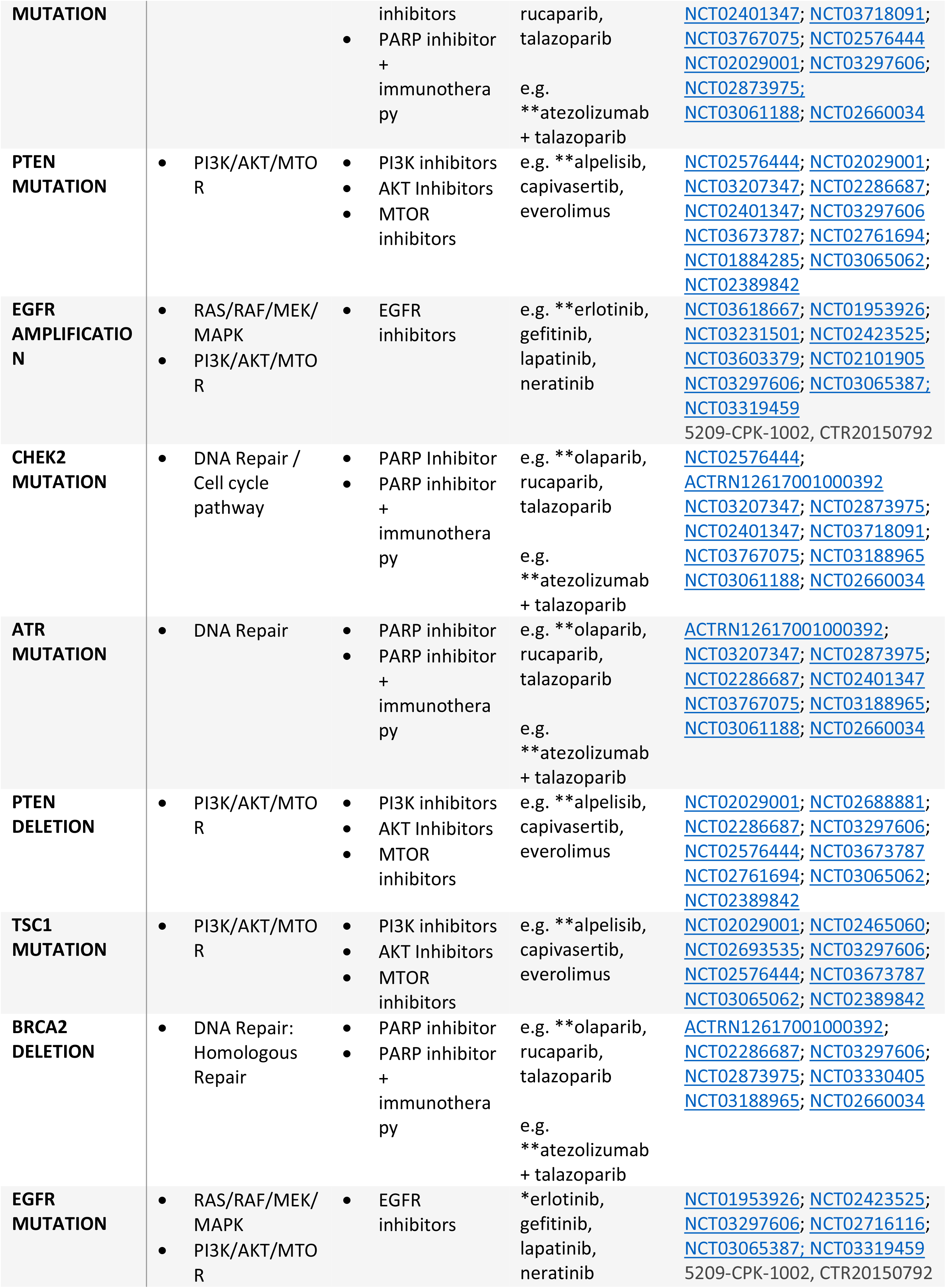
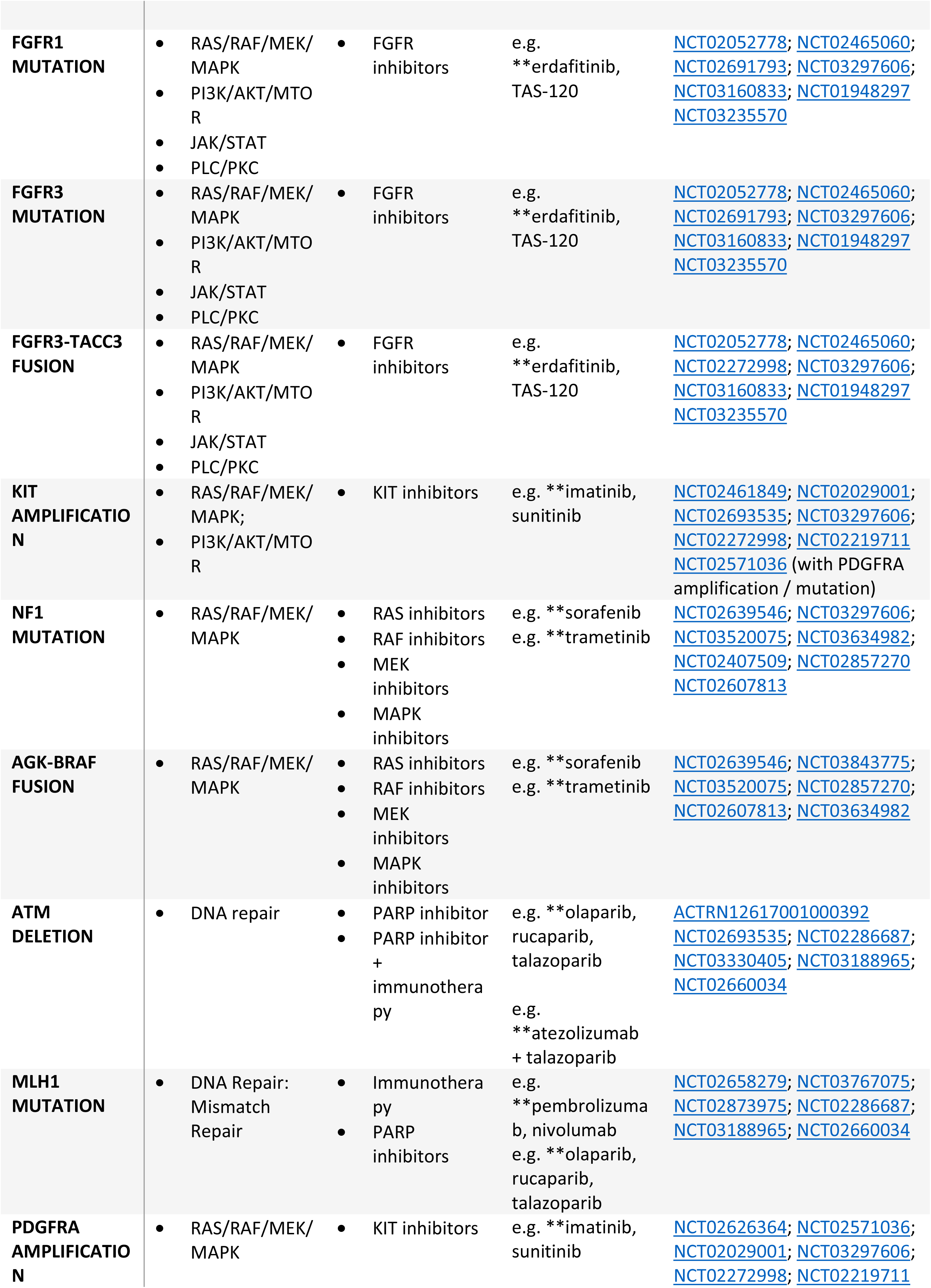
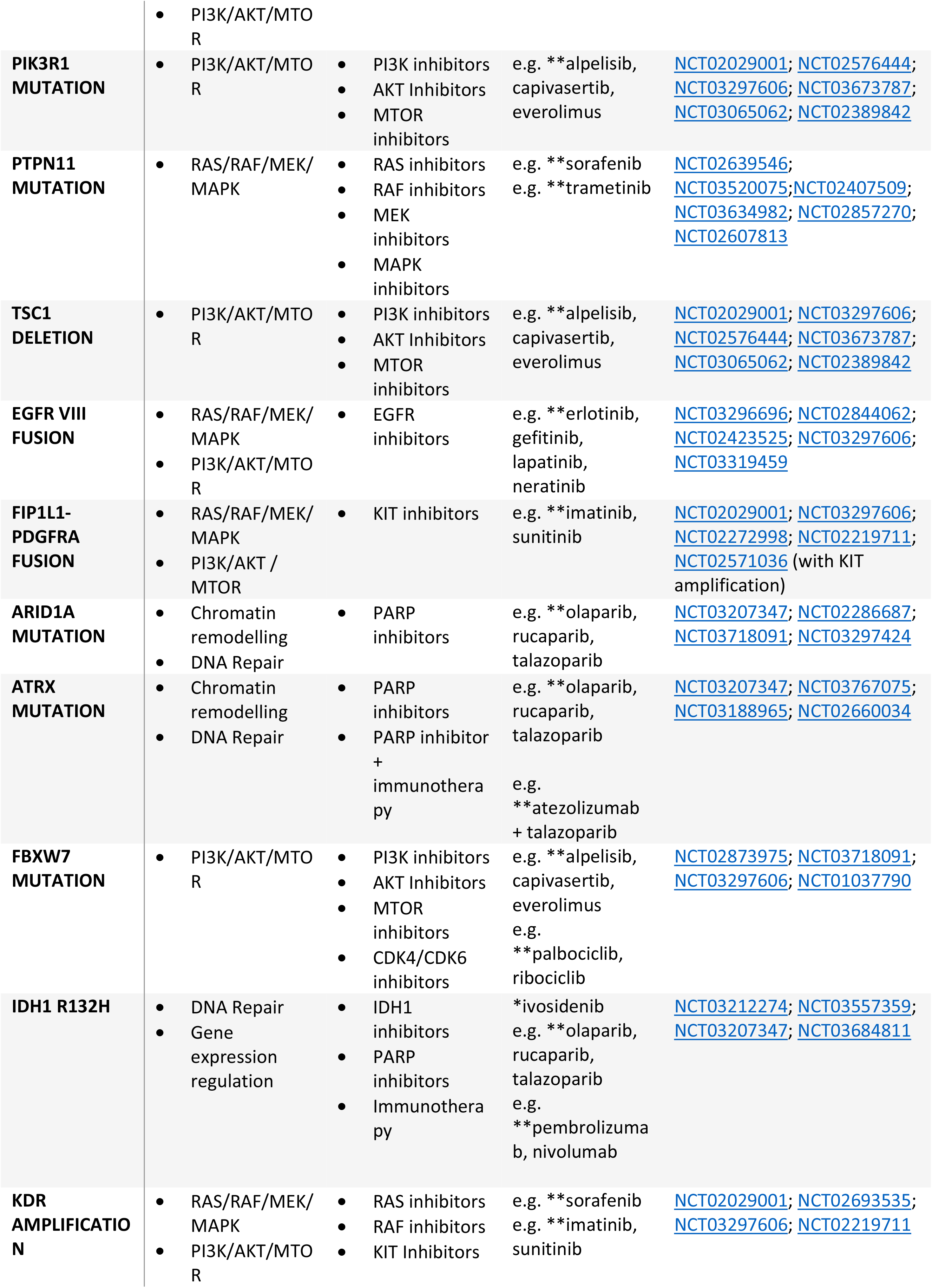
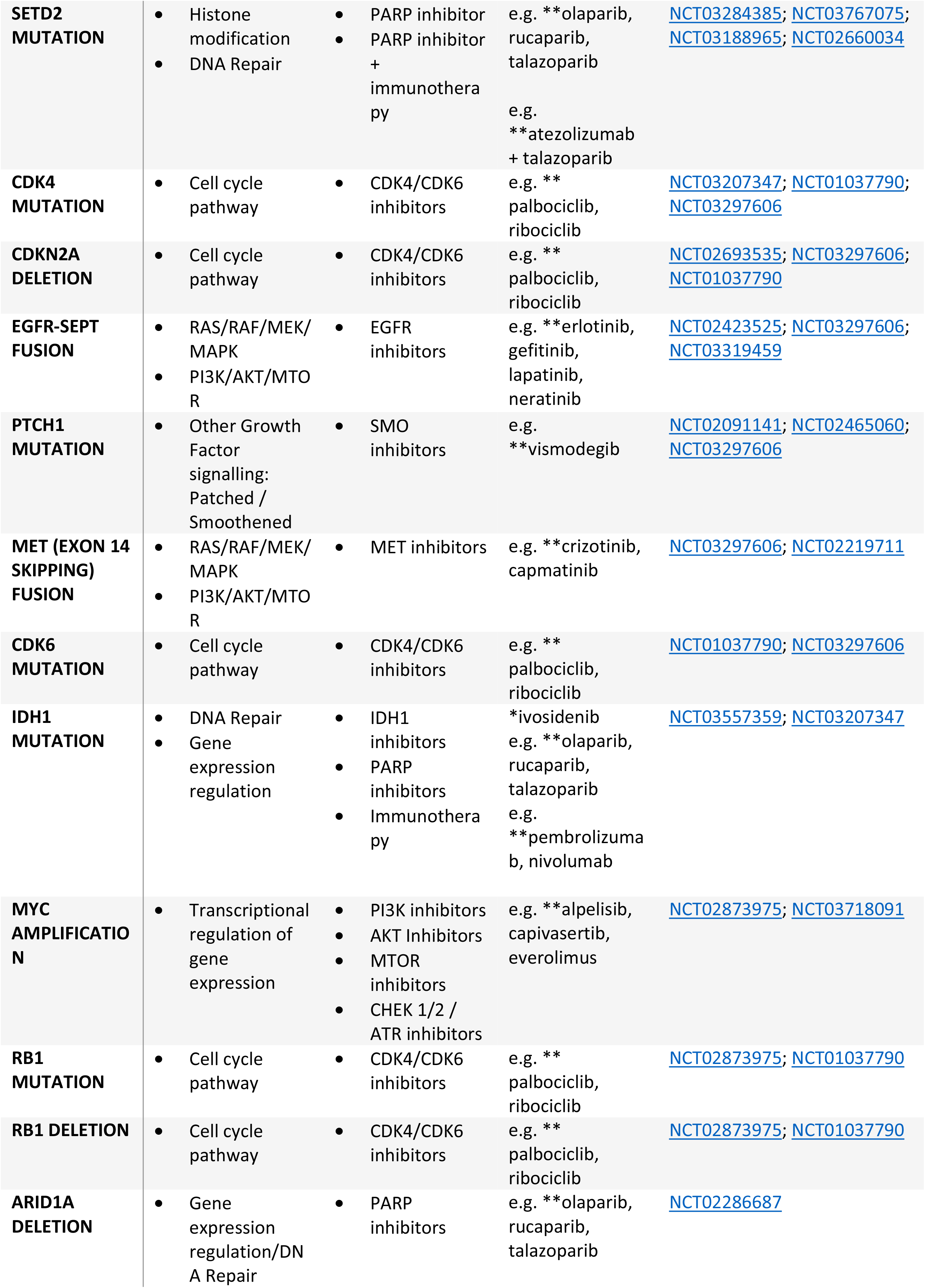
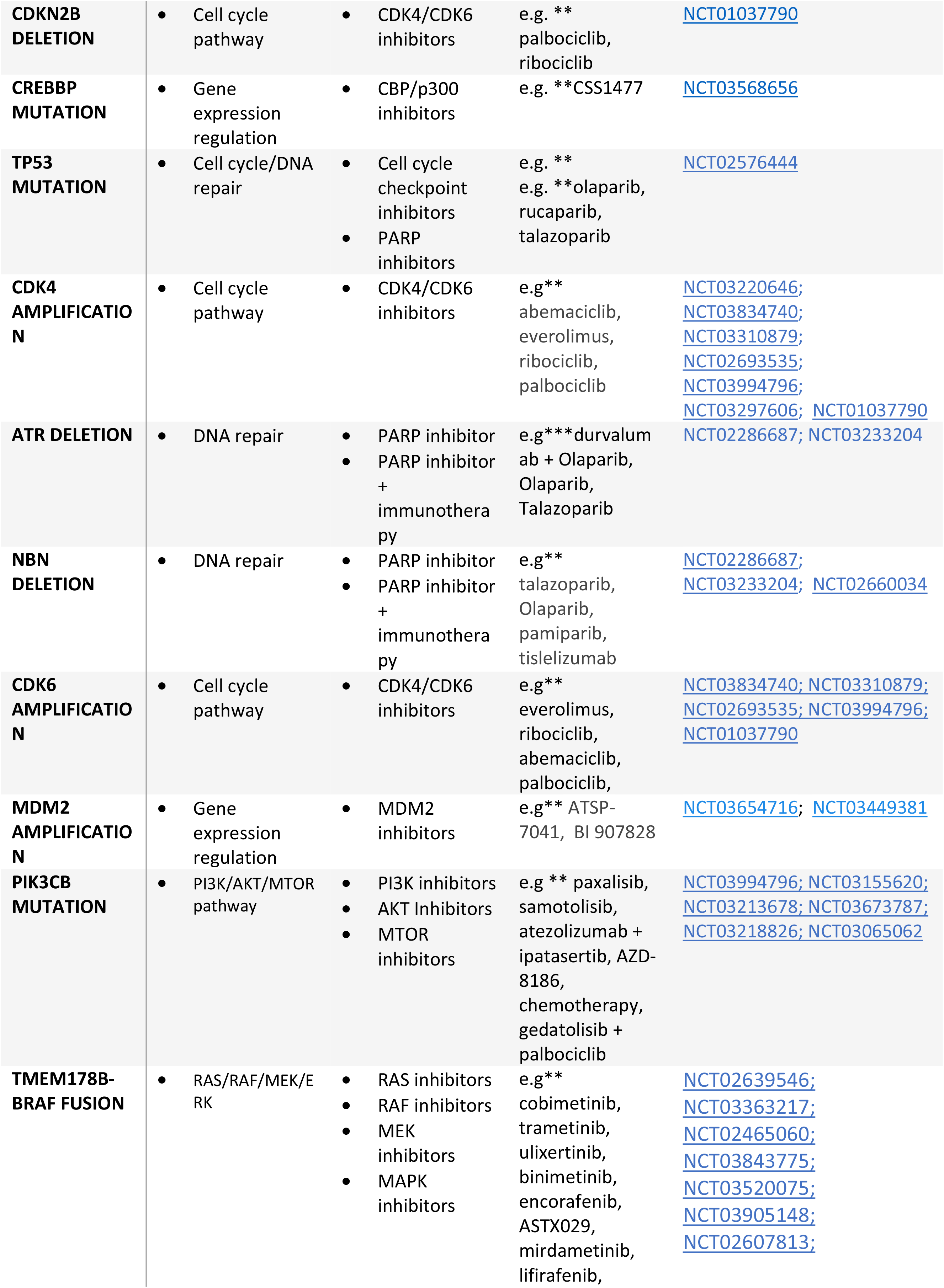
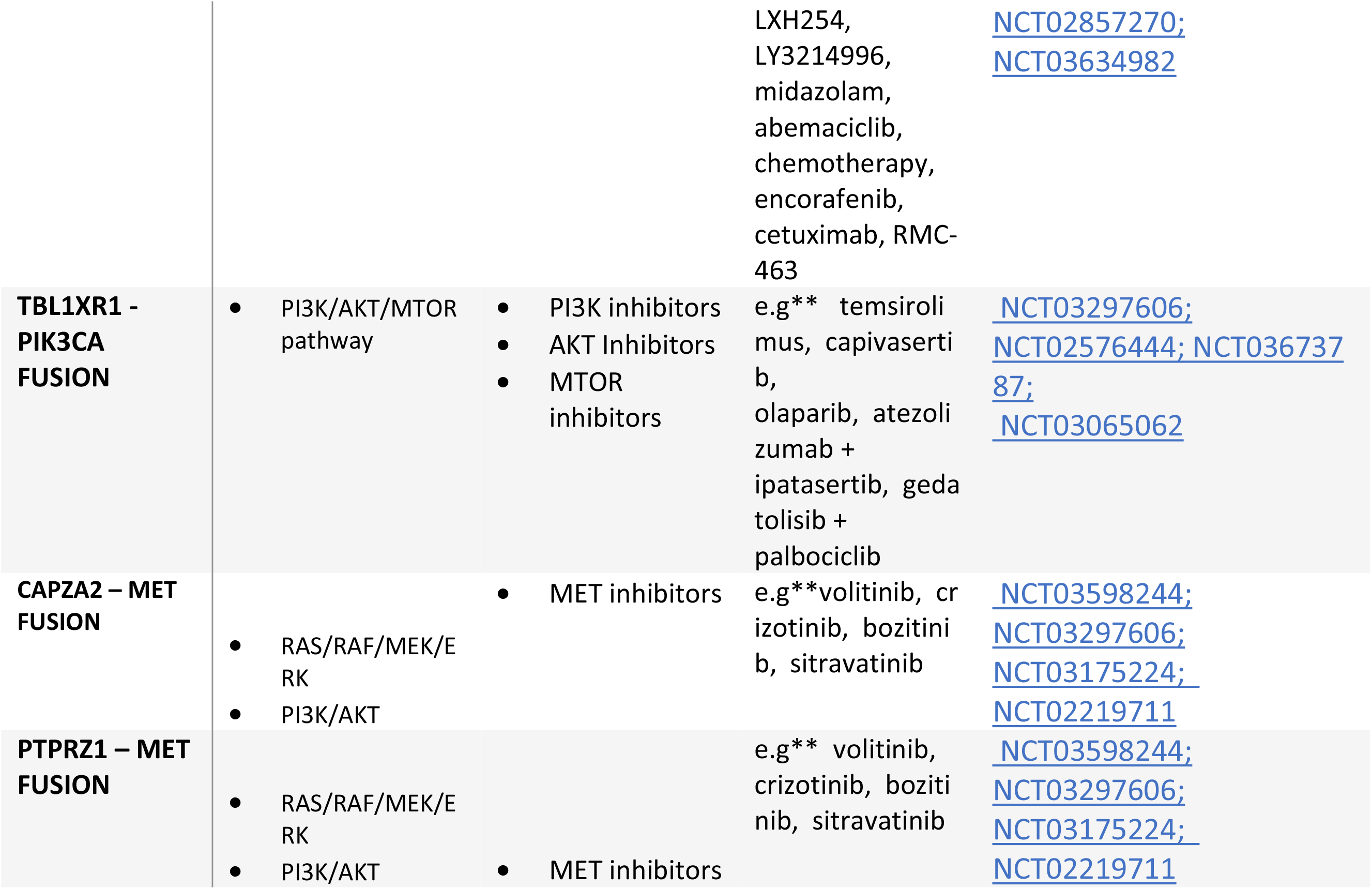
Actionable drug targets in glioblastoma.

Increased levels of PD-L1 tumor cell expression were also identified in a significant proportion of glioblastoma cases. Using a defined predictive cut point of >10% for tumor proportion score, identified as a cut-point for PD-L1 IHC companion diagnostic assays in relation to anti-PD-L1/anti PD-1 directed therapies (21), PD-L1 expression levels were significantly raised in 27.3% of glioblastomas with 12.7% of cases having a tumor proportion score of >50% (S3 Fig). Moreover, actionable variants were identified in 43.6% of patients with aberrations in the DNA repair genes BRCA2 (3.6%), ATM (3.6%), NBN (3.6%), ATR (3.6%), MLH1 (1.8%), ATRX (7.3%), SETD2 (3.6%), PTEN (21.8%), and IDH1 (14.5%). These DNA repair genes share in common evidence based linkage to anti-PD-L1/PD-1 immunotherapies including pembrolizumab, nivolumab, atezolizumab, avelumab, durvalumab and tislelizumab and PARP inhibitors (Table 1). The total percentage of glioblastoma patients with either DNA repair defects or elevated PD-L1 levels amounted to 63.6% of cases and 7.3% of cases were associated with both a DNA repair defect and elevated PD-L1 levels.

## Discussion

Advances in our understanding of somatic cancer genetics and immuno-oncology are now being rapidly translated into clinical practice particularly for advanced metastatic solid tumors. There is now a shift in treatment paradigm from the relatively non-specific empirically directed cytotoxic chemotherapies to a more biologically informed approach where oncogenic somatic mutations are matched with the appropriate targeted agents (22–24). The drug-target pairing that links a dysregulated molecular pathway with a cognate therapeutic agent defines the modern era of precision oncology. This approach has demonstrated superior response rates as compared to nonselective chemotherapy in many tumor types and is associated with less toxicity (12, 13).

Analysis of actionable genomic profiling trending data has revealed unique insights into the complex actional mutational landscape of glioblastoma, the underlying pathobiology that drives this aggressive tumor phenotype and the wide range of potential targeted treatment options available. Actionable variants were identified in all glioblastoma patients tested and indeed the majority of patients harbored three or more actionable variants. This compares to our analysis of solid tumors overall where actionable variants were detected in 90% of patients with a median variant frequency of two (median 2, range 0-13) (25). The genetic variants identified affect many key cancer related regulatory networks including the PI3K/AKT/MTOR, RAS/RAF/MEK/MAPK, JAK/STAT, PLC/PKC signaling pathways, DNA damage repair (DDR) pathways and cell cycle and immune checkpoints. These are key signaling networks mediating fundamental processes including cell proliferation, differentiation, cell migration, DNA damage response and apoptosis which are involved in homeostasis across all tissue types (26–28). This may account for the fact that the majority of detected actionable genomic variants are independent of tumor type or tissue of origin. The high frequency and the presence of multiple actionable mutations identified for each patient tested highlights the broad range of potential personalized treatment options that are available for the treatment of this aggressive disease (Table 1). Many of the targeted agents are directed against the aberrant oncoprotein directly, for example erdafitinib against mutated FGFR1 and FGFR3 and imatinib and sunitinib targeting KIT amplification. Alternative treatment protocols have also been developed targeting components of the cognate signaling pathways downstream. For example, the TBL1XR1-PIK3CA fusion is targeted with therapies inhibiting PI3K directly (alpelisib) or alternatively using inhibitors downstream including AKT (capivasertib), MTOR (temsirolimus) and CDK4/6 (abemaciclib) (Table 1).

Although actionable fusion genes are a relatively rare occurrence in most tumor types (20, 29), here we show that glioblastoma harbors a high frequency of actionable oncogenic fusion genes, in 23.6% of cases. This is in keeping with the observation that mRNA fusions transcripts, including non-druggable fusions, have been identified at high frequency (65%) in glioblastoma (30) Notably all fusions share in common interaction with one or more of the major oncogenic signaling pathways namely RAS/RAF/MEK/MAPK, PI3K/AKT/MTOR,JAK/STAT or PLC/PKC signaling cascades (Table 1). This includes the two novel fusions genes we have identified in glioblastoma, TBL1XR1-PIK3CA and FIP1L1-PDGFRA. Interestingly, the mean number of actionable mutations was greater in samples containing fusions or deletions. This may relate to the fact that fusions and deletions are an indicator of a high degree of cancer cell genome instability (20, 31). Moreover fusion tyrosine kinases have been shown to compromise the fidelity of DNA repair mechanisms which can lead to the accumulation of additional genetic aberrations thus functioning as “first hit” oncogenic aberrations (32–34).

A wide range of DNA repair genes were also found to be mutated in this cohort of glioblastoma patients including ATR, ATM, ATRX, IDH1, BRCA2, NBN, PTEN, SETD2 and the MMR related gene MLH1. These aberrations were bioinformatically linked to either monotherapy with PARP inhibitors harnessing a synthetic lethality approach to treatment or alternatively anti-PD-L1/PD-1/CTLA-4 directed immunotherapy or in combination (Table 1). The linkage with immunotherapy is a consequence that deleterious mutations in DNA repair genes can give rise to an increase in neoantigens (35). Neoantigens in turn are able to elicit tumor-specific immune responses which can then be amplified by the immune-activating actions of immunotherapy including inhibitors of CTLA-4, PD-1 and PD-L1. This linkage is highlighted in the recent landmark FDA approval for PD-1 inhibitors in MMR-deficient tumors (36). Combination therapy with PARP inhibitors and immunotherapy is a compelling strategy and early trials utilizing this strategy are showing encouraging results (37).

Around 8-23% of glioblastoma patients show response to anti-PD-L1/PD-1 directed immunotherapies and the challenge now is to identify more effective companion diagnostic markers that can further refine identification of this treatment responsive group. (38). In keeping with previous studies we observed varying levels of upregulation of PD-L1 in a high proportion of (62%) of cases (39). PD-L1 expression levels have been identified as a predictor of response in some tumor types but is a relatively poor biomarker of response (40, 41). Interestingly a subpopulation of cases in our cohort (7%) showed both elevated PD-L1 expression levels and aberration of DNA repair genes and we are currently investigating whether this signature can be used as a potential improved predictor of immune response across all tumor types (42).

Notwithstanding the high frequency of actionable variants observed in our cohort, EGFR amplification was detected at the lower end frequency to that previously reported. The frequency of reported EGFR amplification is highly variable (8-50%) and is influenced by many parameters including ethnicity and glioblastoma subtype (30, 43, 44). The relatively lower frequency observed in our cohort may be a consequence of the fact that samples were sourced globally from different ethnic populations and also included secondary glioblastomas in which EGFR amplification events are a rare event (43, 44).

In summary we have shown that comprehensive semiconductor precision oncology profiling for actionable variants can significantly increase the potential therapeutic armamentarium available for the treatment of glioblastoma compared to the biomarkers currently used in routine clinical practice and notably targeted therapies in clinical trials are showing promising results (45–51). Interestingly, the majority of actionable genomic variants identified were not tumor type specific, reinforcing the “site agnostic” approach to genomic profiling and supporting the concept of “molecular basket” clinical trials (52). The adoption of semiconductor sequencing methodologies enables clinically directed targeted precision oncology profiling to be applied robustly to routine FFPE clinical biopsy samples allowing integration with established routine diagnostic pathology workflows.

## Author contributions

ML and GW were responsible for study design, conceptualization, supervision and manuscript preparation. RT and KH performed the data analysis and visualization.

## Data availability

The datasets analysed during the current study are provided in the Supplementary File. Further technical details are available from the corresponding author upon reasonable request.

## Ethical statement

The research conducted in this study was limited to secondary use of information previously collected in the course of normal care (without an intention to use it for research at the time of collection) and therefore does not require REC review. The patients and service users were not identifiable to the research team carrying out trend data analysis. This is also in accordance with guidance issued by the National Research Ethics Service, Health Research Authority, NHS and follows the tenants of the Declaration of Helsinki.

## Acknowledgements

The authors thank the technical support staff at Oncologica UK Ltd for their valuable contribution to the study.

